# ShapeUpLMD: An Automated Pipeline for Spatially Optimized Laser Microdissection in Multi-omic Tissue Profiling

**DOI:** 10.64898/2025.12.19.695433

**Authors:** Joshua P. Schaaf, Dave Mitchell, Jonathan D. Ogata, Sakiyah A. TaQee, Katlin N. Wilson, Kelly A. Conrads, Jeremy Loffredo, Brian L. Hood, Tamara Abulez, Allison L. Hunt, Andrea Langeland, Paulette Mhawech-Fauceglia, Kathleen M. Darcy, Christopher M. Tarney, G. Larry Maxwell, Thomas P. Conrads, Nicholas W. Bateman

## Abstract

Laser microdissection (LMD) enables enrichment of defined cellular populations from heterogeneous tissues, providing histologically-resolved molecular profiles with spatial resolution. However, the absence of standardized methods for generating optimized regions of interest (ROIs) limits reproducibility and scalability for spatial multi-omic workflows. To address this gap, we developed ShapeUpLMD, an open-source software tool and integrated workflow that automates the optimization of spatially defined ROIs for LMD, directly from digital pathology annotations.

Fresh-frozen uterine serous carcinoma (USC) tumors (n = 11) were sectioned onto polyethylene naphthalate slides, scanned and subsequently annotated by an expert pathologist. Tumor and non-tumor ROIs were used to train tumor-specific classifiers in HALO (Indica Labs). Classified tumor or unbiased whole tissue ROIs were refined and optimized by ShapeUpLMD prior to automated collection on a Leica LMD7 microscope. Across the cohort, predicted tumor ROIs increased effective tumor purity by 86 ± 6% relative to whole tissue while maintaining high spatial concordance between predicted and collected ROIs (accuracy = 0.9 ± 0.05). Data-independent acquisition mass spectrometry quantified upwards of ∼6,500 proteins across spatially resolved ROIs, revealing regionally coherent clustering of adjacent tumor regions and abundance patterns consistent with tumor and non-tumor cell admixture in an unbiased spatial proteomic application.

ShapeUpLMD provides an automated, reproducible, and scalable framework that bridges digital pathology with LMD enabling high-fidelity spatial enrichment for multi-omic analyses. This workflow increases throughput, reduces inter-operator variability, and supports standardized, regionally resolved tissue collection for spatial systems biology applications. The software is available at https://github.com/GYNCOE/ShapeUpLMD.

## Introduction

Laser microdissection (LMD) enables the precise isolation of specific cellular populations from histologically complex tissues, facilitating high-resolution spatial molecular analyses of tissue microenvironments. This technique mirrors workflows used in diagnostic pathology, where tissue sections are mounted onto slides, stained, and cellular architecture is visually assessed by expert pathology review to identify distinct cellular architectures or regions of interest (ROIs). By adding this spatial axis to molecular analyses, LMD serves as a powerful bridge between histopathological context and downstream molecular characterization.

Emerging evidence has shown that cells collected by LMD yield high quality molecular analytes including DNA, RNA and protein from solid tumor samples (1). This technique has proven especially valuable for microcompartment analysis of tumor tissues, which often exhibit marked heterogeneity arising from admixture of malignant, stromal, and immune cell populations (1,2). By enriching histologically homogeneous subpopulations, LMD enables disentanglement of molecular signals that would otherwise be obscured in bulk tissue analyses. More recently, LMD has become integral to spatial -omics workflows, such as deep visual proteomics (3), supporting high resolution characterization of molecular heterogeneity within tumor architectures. As LMD adoption increases in high-throughput multi-omic workflows, standardized collection protocols and quality control practices will be paramount to ensure reproducibility, scalability, and intra- and inter-study comparability.

Traditionally, ROI selection for LMD enrichment relies on manual annotation by expert pathologists - a time-intensive process subject to inter-individual variation. Manual annotation directly on the LMD microscope also exposes tissues to ambient conditions for extended periods, potentially compromising molecular integrity. Recent advances have enabled direct transfer of digital tissue ROI annotations onto LMD instruments, supporting more efficient and automated collection of defined cell populations (4–6). Innovations in digital pathology and machine learning-based image analysis present new opportunities to automate ROI identification, thereby improving reproducibility and throughput. Current workflows lack strategies to algorithmically optimize ROI geometry, such as standardizing ROI shape and size, controlling inter-ROI spacing, and defining regionally resolved sampling patterns, to fully support spatial multi-omic analyses. Consequently, there remains a critical need for standardized and automated methods that generate optimized, spatially defined ROIs to minimize inter-operator variability and maximize reproducibility of collections at cohort-scale.

To address this need, we developed ShapeUpLMD, an open-source software tool that automates generation of spatially defined, LMD-optimized ROIs from digital pathology annotations. ShapeUpLMD extends our prior efforts integrating digital pathology with automated LMD (4) by introducing algorithms for ROI refinement, area and spacing optimization, and spatial partitioning to support spatially resolved tissue sampling. We demonstrate performance of this pipeline in a cohort of uterine serous carcinoma (USC) tumors and illustrate its application for spatial proteomics with a representative USC tissue specimen. Together, these efforts establish a scalable and reproducible pipeline for automated spatial enrichment in multi-omic workflows.

## Methods

### Patient Tissues and Classification of Tumor Regions of Interest

Fresh-frozen tumor tissues were collected from eleven USC patients enrolled in a WCG IRB approved #20110222 Tissue and Data Acquisition Study of Gynecologic Disease who underwent surgery at Inova Fairfax Medical Campus (Falls Church, VA, USA), University of Oklahoma Health Sciences (Oklahoma City, OK, USA) or at University of Virginia (Charlottesville, VA, USA); all experimental protocols involving human data in this study were in managed in accordance with the Declaration of Helsinki. Thin (8 µm) tissue sections were sectioned and placed on glass or polyethylene naphthalate membrane slides (PEN, Leica Microsystems, Wetzlar, Germany) into which fiducials were pre-cut for image co-registration as previously described (4). After staining with aqueous hematoxylin and eosin, tissue sections were scanned (Aperio ScanScope XT slide scanner, Leica Microsystems, Wetzlar, Germany) and digital images were reviewed by a board-certified pathologist (P.M.F.) to confirm disease diagnosis and define representative ROIs.

Representative tumor and adjacent non-tumor regions were annotated on adjacent PEN membrane tissue sections and HALO Image Analysis Platform v4.0.5107 (Indica Labs, Albuquerque, New Mexico, USA) was used to train tumor cell-specific classifiers using the DenseNet 2 module. A minimum of ten high-purity tumor regions were manually annotated on a PEN slide image, guided by annotations from expert pathology review of adjacent H&E tissue sections (please see Supplementary Figure 1 for more details). To ensure spatial and morphological diversity, five regions were placed near the periphery of the tissue section area, while the remainder were distributed more centrally when possible. Tumor shapes were then positioned adjacent to non-tumor elements or background glass to define edge boundaries for model training. A separate annotation layer was created in which ten non-tumor annotations were added, each encompassing a tumor annotation region noted above to maximize differentiation of tumor from non-tumor regions. Non-tumor annotations were also chosen to include stromal elements and background glass, taking care to ensure that each non-tumor regions contained only the embedded tumor annotation as the sole source of tumor tissue. These annotations were used to create cell type-specific classifiers using the DenseNet 2 image analysis module. Minimal object size was set to 10,000 µm² to exclude small, irrelevant shapes, and the resolution was set to 1 µm/pixel to allow for tighter classification boundaries. Two classes were defined, tumor and non-tumor, with tumor set as the priority class to ensure more conservative inclusion of ambiguous regions. Training was run for approximately 3,000 iterations and the resulting high-resolution binary tumor cell-specific classifier was used for downstream analysis.

### ShapeUpLMD – Development and Application

ShapeUpLMD was developed in Python v3.10.16. Regions determined to be collected were retrieved from the HALO Image Analysis Platform, generated using HALO AI v4.0.5107 (Indica Labs, Inc.) and the HALO GraphQL database. Annotations were converted from vertex coordinates into shapely v2.0.6 polygon objects, making sure to account for all HALO shape types, and whether that shape is to be collected or excluded (isExclusionRegion). Exclusion regions were connected to their encompassing collected shape edge to avoid collecting negative regions due to LMDs gravitational collection. The shortest path from the exclusion region to the encompassing shape edge was used to diminish the loss of collection yield. Shapes were then reduced in size through iteratively suggesting horizontal and vertical slices, choosing the proposed sliced sections which provide the largest area of collectable material. For this application, we defined maximum shape area to be 0.36 mm^2^ and slice width as 60 µm (as applied for tumor-enriched collections), both set to avoid tissue peeling during cutting. Regions between shapes were widened so that they were at least 60 µm apart. This may introduce thin shapes, so shapes which had thin regions of less than 60 µm wide were removed recursively. Shapes were finally simplified with the simplify function in shapely using the Douglas-Peucker algorithm, as we identified that fewer vertices enabled faster cutting times on Leica’s LMD7. Shapes smaller than 0.01 mm^2^ were removed between multiple ShapeUpLMD steps, as they were deemed too small for collection. Shapes could then be placed into spatially resolved groups by their proximity to algorithmically determined points spread evenly within the tissue section. Finalized shapes were exported to an LMD7-importable extensible markup language XML file using python code modified from the Malleator tool (4). Slide image metadata such as micrometers per pixel and slide dimensions were read using tiatoolbox (1.6.0). Environment files, HALO database connection files, and ShapeUpLMD code are available on GitHub (https://github.com/GYNCOE/ShapeUpLMD).

### Collection of ROI shapes

Tumor regions annotated in HALO were optimized for LMD using ShapeUpLMD. Briefly, tumor layer coordinates were accessed through the HALO API and subdivided into uniformly spaced polygons using the following parameters; 60 µm spacing between shapes, with minimum and maximum dimensions of 100 × 100 µm and 600 × 600 µm, respectively. For unbiased collections, we kept the same minimum size but updated the spacing to 50 µm and reduced the maximum size to 250 × 250 µm. These constraints were implemented to minimize membrane and tissue curling while ensuring adequate sample areas for recovery from collections using a 5× objective. For spatial analyses of tumor-enriched regions, each subdivided region was assigned a spatial cluster identifier using the spatial_segmentation_brute_force function in ShapeUpLMD, and total cluster areas (mm²) were balanced across the slide. For unbiased spatial analyses, spatial regions were similarly generated, but by using rebalance_multipolygons_by_area function in ShapeUPLMD to achieve more balanced area sizes across the tissue section of interest. Cluster-specific XML annotation files were generated and imported into the Leica LMD7 system. Alignment was verified using fiducial markers to maintain spatial correspondence with the tissue section as previously described (4) and ROIs were cut using the following LMD microscope software settings: power 60, aperture 2, speed 33, specimen balance 5, line spacing (draw + scan): 5, head current 100 %, pulse frequency 555 and offset 45. LMD tissue collections from each spatial cluster were collected separately into pressure cycling technology (PCT) tubes (please see Sample preparation for quantitative proteomic analysis below for more details).

### Evaluation of Manual Tumor, Automated Tumor, and Collected Regions

Tumor cell-specific ROI annotations generated by manual inspection, by automated classifier generation ROIs as noted above, by ShapeUpLMD-derived and optimized ROIs, and finally collected ROIs were all compared using a series of in-house tools generated in Python. Briefly, whole slide image annotation polygons corresponding to ground truth tumor, predicted tumor, and tissue regions were converted into binary masks using custom functions utilizing the packages NumPy, Shapely, and OpenCV. All performance metrics were computed within tissue boundaries; pixel-wise comparisons between ground truth and predicted tumor masks were used to calculate precision (the proportion of predicted tumor pixels that were correctly identified), recall or sensitivity (the fraction of true tumor pixels correctly detected), F1 score (the harmonic mean of precision and recall summarizing segmentation balance), and accuracy (the overall proportion of correctly classified pixels). Collected regions were estimated by the HALO AI DenseNet 2 model which determines areas collected from post-LMD collection scanned slides. Slides were loaded using OpenSlide, and code was created to process and generate color-coded overlay masks, thumbnail previews, and a timestamped CSV file summarizing all metric results for subsequent visualization and statistical aggregation. Mann Whitney U testing was performed using MedCalc v20.109.

### Sample preparation for quantitative proteomic analysis

Tumor regions of interest (ROIs) were imported onto an laser microdissection microscope (LMD7, Leica Microsystems, Wetzlar, Germany) from XML coordinate exports and co-registered with mounted slide images as previously described (4). Tumor enriched ROIs were collected by laser microdissection into 20 µL of 100 mM triethylammonium bicarbonate (TEAB)/10% acetonitrile (ACN), pH 8.0 and unbiased ROIs were collected in 100mM ammonium bicarbonate/10% ACN, in MicroTubes (Pressure BioSciences, Inc, South Easton, MA) followed by digestion with a heat-stable form of trypsin (SMART Trypsin, ThermoFisher Scientific, Inc.) using pressure cycling technology with a barocycler (2320EXT Pressure BioSciences, Inc). Digested peptide concentrations were determined using the bicinchoninic acid assay (BCA assay, ThermoFisher Scientific, Inc.) for tumor-enriched collections and equivalent volumes of peptide digest were analyzed for tumor-enriched and unbiased ROI collections.

### Liquid-Chromatography-Mass Spectrometry (LC-MS) Analysis

For tumor enriched analyses, a timsTOF Ultra 2 MS connected to an EvoSep LC system (EvoSep, Odense, Denmark) via a CaptiveSpray 2 source (Bruker, Billerica, MA) was used for dia-PASEF LC-MS/MS analysis. Peptide digests were separated using a C18 IonOpticks column (particle size 1.7 μm, 75 μM μm ID, 15 cm length) at a flow rate of 0.2 μL/min; solvent A (0.1% formic acid in water), solvent B (0.1% formic acid in acetonitrile), using the Evosep Whisper Zoom 40 SPD method. For dia-PASEF analysis, the scan range was *m/z* 400-1000 with a mobility range of 0.64 to 1.37 V·s/cm² with eight MS/MS ramps per MS scan. Three MS/MS scans were collected per ramp starting at *m/z* 400, 600, and 800 with a width of 25 which increased by 25 Th for each successive ramp. The capillary voltage was set at 1600 V, and the ramping collision energy was 20 eV at 0.6 mobility to 59 eV at 1.6 mobility. dia-PASEF scan range *m/z* 100–1700 in positive mode, and IMS service ramp time of 100 ms.

For unbiased spatial proteomic analysis, an Orbitrap Astral (ThermoFisher Scientific, Inc) MS was connected to a Vanquish Neo (ThermoFisher Scientific, Inc) using an Easy-Spray source (ThermoFisher Scientific, Inc.). Peptides were loaded onto an Acclaim PepMap 100 pre-column (75 μM μm ID x 2 cm) and separated on an IonOpticks Elite XT analytical column (1.7 μm, 75 μM μm ID x 15 cm) at a flow rate of 0.35 μL/min; solvent A (0.1% formic acid in water), solvent B (0.1% formic acid in acetonitrile) from 0-25 %B over 25 min, 25-35 %B over 5 min, 35-99 %B over 5 min followed by column washing at 99 %B for 6.3 min before equilibrating (5 column volumes) for the next injection. For data-independent (DIA) analyses, full MS data were collected in profile mode from *m/z* 380-980 at a resolution of 240k. Voltage was set to 1900; normalized AGC target was 500% (5e6 absolute), RF Lens was 40%, with a maximum injection time of 100 ms using Advanced Peak Determination and lock mass correction (*m/*z 445.12003; 10 ppm mass tolerance). DIA data were collected in the Astral detector over the same precursor mass range with isolation windows of 2 Th and maximum injection time of 3 ms. The normalized AGC target was 500% (5e4 absolute), HCD collision energy was 27, an RF Lens of 45% and a scan range of *m/*z 150-2000 using a loop control time of 0.5 s.

### Proteomic Data Analysis

Mass spectrometry RAW files were searched with Spectronaut v19.9 (Biognosys AG, Switzerland) in directDIA mode against a UniProtKB/Swiss-Prot database (Homo sapiens, downloaded May 24, 2024). Default Spectronaut settings were used, except minimum peptide length was set to 6, methionine oxidation was the only variable modification, and precursors passing FDR < 0.01 were considered in downstream analysis. Spectronaut parquet report outputs were imported into R v4.2.33 via the arrow package v16.1.04. The columns Stripped.Sequence, PEP.AllOccuringProteinAccessions, and FG.MS2Quantity were extracted as peptide sequence, mapped protein accessions, and precursor MS2 intensity, respectively. Precursors mapping to multiple proteins were assigned to the protein quantified with the most unique peptides. The sum of sample-level precursor MS2 intensities were normalized to the lowest sum-intensity observed across all samples. Proteins observed in ≤50% of samples were removed and remaining missing values were imputed by k-nearest neighbors (k = 11) via the impute package v1.72.05. Resulting protein abundances were median centered per sample and log_2_ transformed. From the resulting protein-level matrix, tumor-enriched proteome data from 10mm (Fig. 2B) or 5mm (Fig. 2C) regions were subset and standardized by row-wise z-scoring. Global proteomics data generated for unbiased spatial ROIs for a representative USC tumor tissue section underwent protein-wise quantile standardization across log_2_ fold-change ROI abundance ranges to scale between ± 2.0. Standardized protein abundances were then mapped to corresponding XML coordinates and rendered into tissue section-level proteome abundance maps using in-house tools that included R v4.4.0 and ggplot2 (3.5.2). Tumor and admixture (average of stroma and immune scores) enrichment for unbiased proteome data was assessed using the ProteoMixture tool as previously described (7). Hierarchical clustering was performed with pheatmap v1.0.126 using Canberra distance and Ward.D linkage. Differential analyses of proteins in whole tumor collections and averaged protein abundances across enriched tumor collections was performed using a z-test and proteins significantly altered (z-test p-value < 0.05, FC ± 2.0) were included in downstream Hallmark and Canonical Pathway analyses (FDR < 0.05) using MSigDB (8).

## Results

### Automated Generation of LMD optimized ROIs with ShapeUpLMD

We developed ShapeUpLMD, an open-source, Python-based tool that generates optimized and spatially defined ROIs for LMD from digital histopathology annotations (Figure 1A). To demonstrate proof of concept, we analyzed fresh-frozen thin sections from 11 USC tissue specimens. Expert pathology review of digitally scanned H&E images confirmed disease diagnosis and provided initial annotations of tumor and non-tumor cell populations. These annotations were manually transferred to digital images of PEN membrane slides pre-cut with fiducials, enabling XML coordinate alignment on LMD microscopes as previously described (4). Tumor and adjacent non-tumor regions were used to train a classifier for each patient tumor section using the HALO digital pathology platform (see Supplementary Figure 1 for additional details). The accuracy of automated ROI masks for tumor cell populations was assessed by direct comparison against manual annotations (Supplementary Figure 2A). Co-registration and direct comparison across eleven patient samples demonstrated high concordance between automated and manual tumor annotations (Figure 1B, average accuracy = 0.87 ± 0.05, Supplementary Table 1). Predicted tumor ROIs also exhibited a significant increase in tumor purity over whole tissue areas (Figure 1C, 86 ± 6%, MWU p <0.05). Annotated regions were exported from HALO as XML coordinate files and used by ShapeUpLMD for ROI optimization. ShapeUpLMD optimizes ROIs with user-defined ROI area constraints (in mm²) and distances between ROI areas, such that collection with LMD is possible (see Methods). Optimized ROIs and fiducial coordinates are then exported as LMD-compatible XML files. Once imported into LMD microscope software, fiducial coordinates facilitate the alignment of optimized ROIs with corresponding PEN-mounted tissue sections. Using this workflow, we harvested ShapeUpLMD-optimized tumor cell ROIs from eleven USC tissue specimens and then compared pre-collection optimized-ROIs with classifier-determined collected regions to assess performance. Post-collection assessment showed high fidelity of tissue capture and boundary alignment (Figure 1D, average Accuracy = 0.9 ± 0.05, Supplementary Figure 2B, Supplementary Table 2).

**Figure 1:**
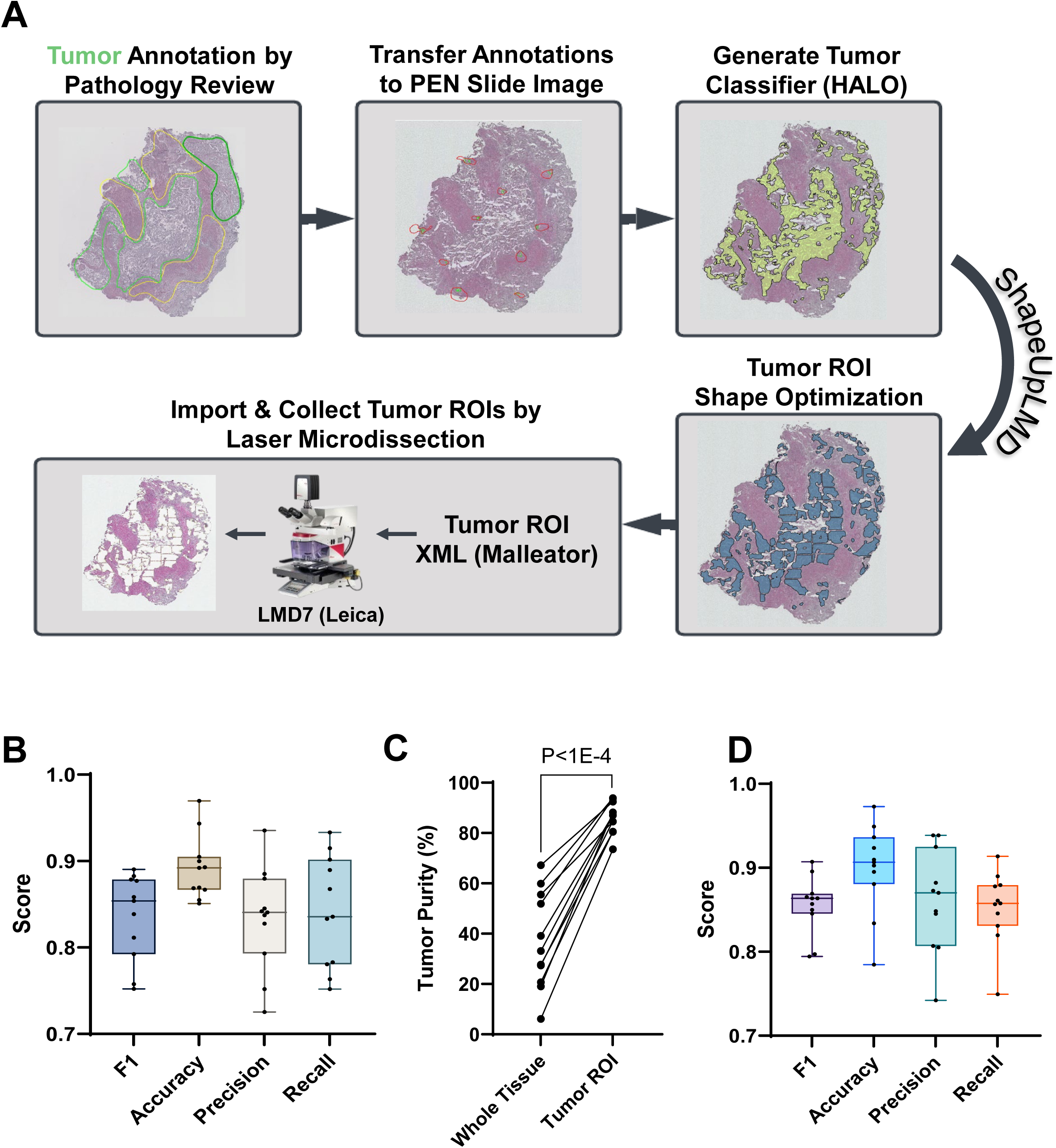
Automated generation, optimization, and laser microdissection of tumor cell regions of interest (ROI) using ShapeUpLMD. **A:** Tumor and non-tumor cell populations are annotated in digital histopathology images by expert pathology review and inform annotation of tumor and non-tumor ROIs on adjacent polyethylene naphthalate (PEN) slide images that include fiducials enabling feature alignment (also See Supplementary Figure 1). Annotated ROIs are used to train a tumor-specific classifier in HALO. Tumor ROIs are then merged with an annotation layer including fiducial coordinates to undergo shape optimization using ShapeUpLMD. Optimized tumor ROIs and fiducial coordinates are then imported onto the laser microdissection microscope for automated sample collection. B: Comparison of manual annotations and automated classifier generated tumor ROIs revealed a high degree of overlap and accuracy between these regions, also with high precision and recall metrics. C: Line plot comparing relative tumor purity in whole slide tissue areas by comparison of automated tumor ROIs to whole tissue areas for 11 USC patient tumors; p-value reflects MWU testing. D: Comparison of automated versus post-collected tumor ROIs revealed a high degree of overlap and accuracy between these regions, also with high precision and recall metrics.

### Spatially-Resolved Tumor ROI Collections with ShapeUpLMD

Once ShapeUpLMD has created collectable regions, ROIs can be split into smaller, spatially defined regions to enable high-resolution spatial analyses. To demonstrate this feature, we subdivided tumor ROIs from a representative USC tissue section into three 10 mm² regions and six 5 mm² regions (Figure 2A). For comparison, we collected whole tissue, enriched tumor ROIs, and spatially defined ROIs of interest. All samples underwent PCT-assisted digestion, and peptides were analyzed by diaPASEF mass spectrometry, resulting in the quantification of 5,294 total proteins across whole tumor, enriched tumor and spatially resolved, ShapeUpLMD-optimized tumor cell ROIs. Unsupervised hierarchical clustering revealed distinct protein expression profiles between whole tumor collections compared to enriched tumor ROIs (Figure 2B & 2C). Spatially defined ROIs clustered according to local proximity, reflecting pronounced regional proteomic heterogeneity within individual tumors (Figure 2B & 2C). Proteins elevated in whole versus tumor specific ROI collections (z-test p-value < 0.05, FC ± 2.0) were enriched for pathways regulating epithelial-mesenchymal transition and extracellular matrix dynamics and included multiple collagen isoforms (COL1A2 & COL3A1) (Supplementary Tables 3 & 4). These findings are consistent with greater stromal cell admixture and lower purity in whole tissue collections, as we have previously reported (2).

**Figure 2:**
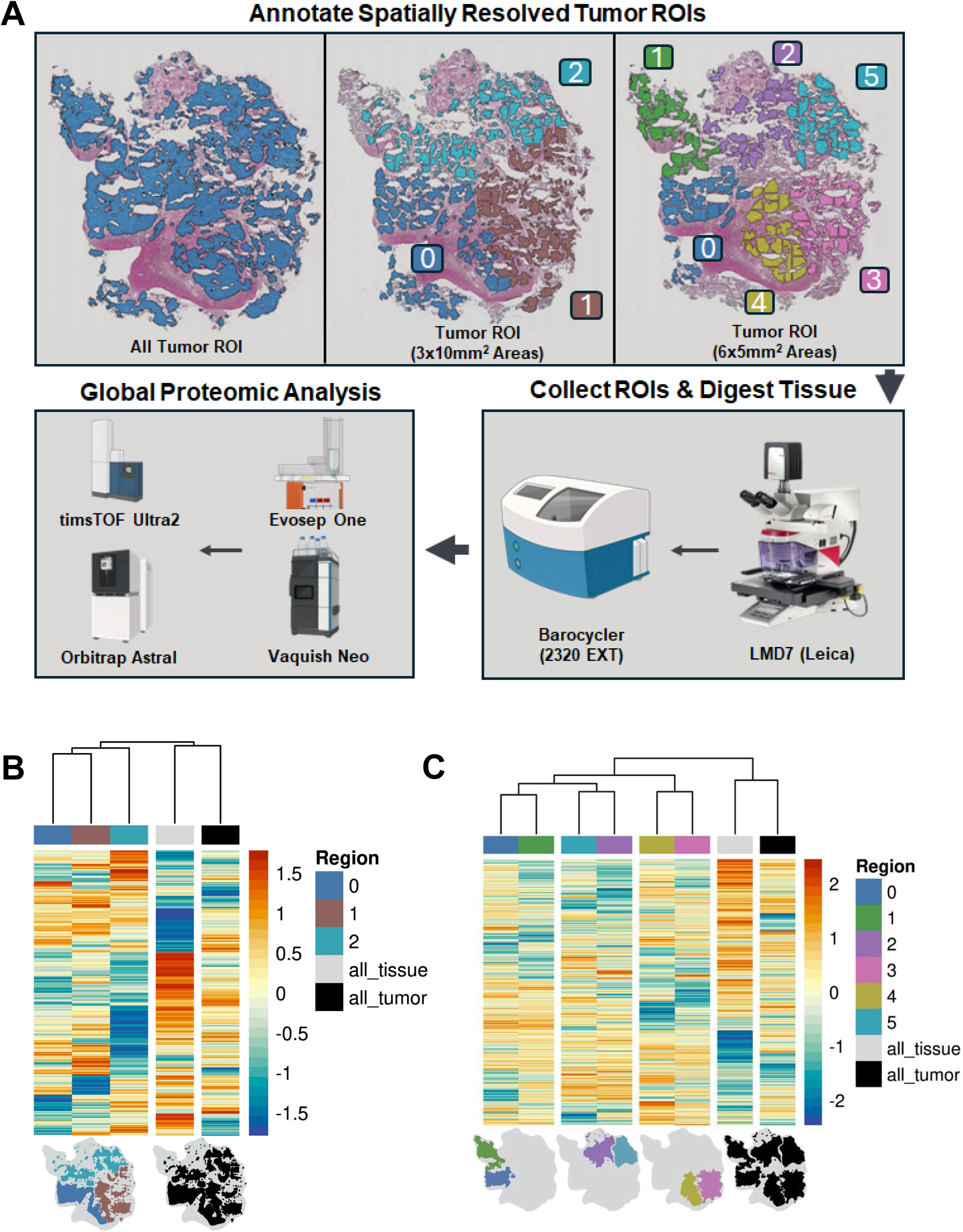
Spatially-resolved proteomics of laser microdissected tumor regions collected using ShapeUpLMD. **A:** Tumor ROIs can be further compartmentalized by the ShapeUpLMD tool into discrete areas to enable spatial collections. Workflow shown here demonstrates annotation of three tumor ROI areas at 10.0 mm^2^ and six tumor ROI areas at 5.0 mm^2^ areas for a single USC tissue sample. Tumor ROIs are then collected by LMD and collected tissues undergo pressure-assisted digestion with trypsin followed by liquid chromatography-data independent acquisition mass spectrometry. B: Unsupervised, hierarchical cluster analysis of proteins quantified across all samples comparing whole (all_tissue), enriched tumor (all_tumor) and three 10.0 mm^2^ tumor ROI areas. C: Unsupervised, hierarchical cluster analysis of proteins quantified across all samples comparing whole (all_tissue), enriched tumor (all_tumor) and six 5.0 mm^2^ tumor ROI areas.

### Spatially-Resolved Unbiased ROI Collections with ShapeUpLMD

To evaluate the utility of ShapeUpLMD for unbiased spatial proteomics, we generated whole tissue equivalent ROIs (∼1 mm² each) spanning the entire tissue area from a representative USC tumor section (Supplementary Figure 3). ShapeUpLMD generated fifty-five uniformly sized ROI that were independently collected by LMD, which were processed and analyzed by DIA mass spectrometry on an Orbitrap Astral mass spectrometer, resulting in 6,536 proteins that were co-quantified across all tissue ROIs.

Tumor, stroma, non-tissue / necrotic tissue regions were predicted using the workflow detailed above (Supplementary Figure 1, Supplementary Table 5). Tumor purity and non-tumor (stroma and immune) cell admixture scores were inferred from global proteome data using ProteoMixture (7) Supplementary Table 6). Classifier-based spatial predictions showed strong concordance with proteome-derived estimates of tumor content (*r* = 0.63, P<1E-4) and non-tumor cell admixture (*r* = 0.76, P<1E-4) ProteoMixture scores, demonstrating that ShapeUpLMD enables spatially-resolved sampling that retains underlying biological composition.

Unsupervised hierarchical clustering of the top most variably abundant proteins revealed that spatially unbiased ROIs segregated according to predicted tumor and non-tumor composition (Figure 3A). Well-established epithelial (e.g., E-cadherin, CDH1 and (epithelial cellular adhesion molecule, EPCAM) and stromal markers (e.g. fibroblast activating protein, FAP and collagen 1A1, COL1A1) exhibited distinct regional localization patterns (2), with elevated abundance profiles in ROIs enriched for their respective cellular compartments (Figure 3B, Supplementary Table 6). These findings resonate prior studies demonstrating proteomic divergence between tumor and non-tumor (stroma and immune) cell populations in gynecologic malignancies (1,2,9,10) and confirm that ShapeUpLMD enables spatially resolved proteome mapping without prior ROI bias.

**Figure 3:**
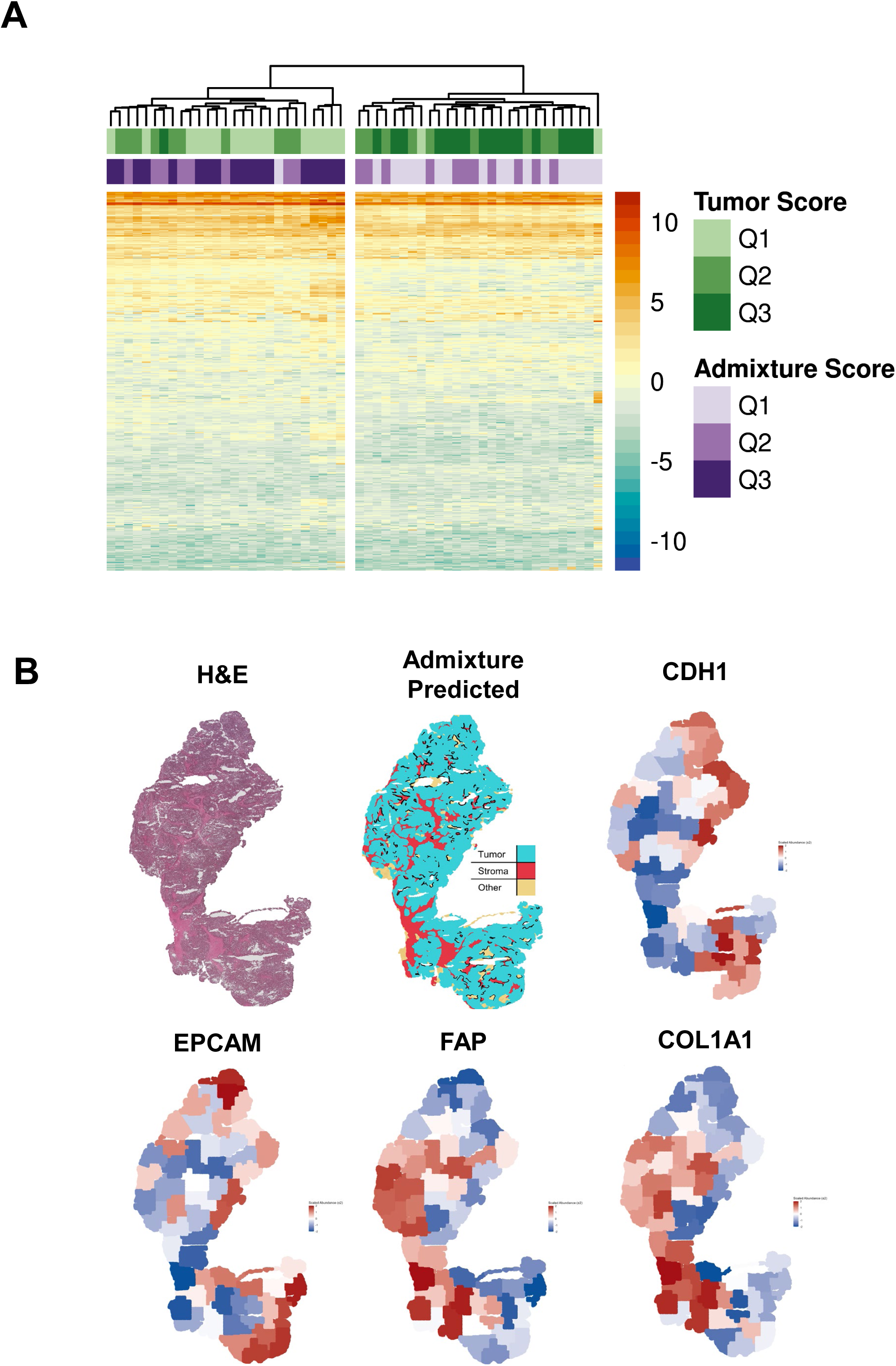
Unbiased spatial proteomic mapping of a uterine serous carcinoma tissue section using ShapeUpLMD. A: Unsupervised hierarchical clusters analysis of the top 1,000 most variably abundant proteins (median absolute deviation) quantified across fifty-five ∼1.0mm^2^ ROIs spanning a single representative tissue section from a uterine serous carcinoma tumor. Metadata reflects predicted tumor or stroma & immune admixture scores calculated using ProteoMixture (7) and is categorized as tertiles, Q3 = increased score, Q1 = decreased score. B: Representative hematoxylin and eosin (H&E) stained tissue section, admixture predicted from automated classifications defined as tumor, stroma and other (necrotic and tissue-free space) ROIs, as well as spatial maps of representative tumor protein markers (E-Cadherin, CDH1 and epithelial cellular adhesion molecule, EPCAM) as well as representative stroma markers (fibroblast activating protein, FAP and collagen 1A1, COL1A1), derived from spatial proteomic analyses.

## Discussion

Recent developments in spatial genomics and proteomics have underscored the importance of precisely and reproducibly characterizing cellular subpopulations in heterogeneous tumor microenvironments. Laser microdissection has emerged as a key enabling technology for this purpose, yet limitations in standardization, reproducibility, and throughput continue to restrict its widespread adoption. A central challenge to scaling LMD lies in the manual generation of regions of interest (ROIs) for enrichment of cellular subpopulations. This process is time-intensive, may expose tissue to ambient conditions for extended periods, and is subject to inter-operator variability.

Our group recently addressed several of these challenges through the development of Malleator (4), a software application that enables direct import of ROI coordinate files generated from digital slide images onto LMD microscopes to support automated tissue collections. Recent methods focused on increasing reproducibility for LMD-based experiments include pyLMD (5) and QuPath-to-LMD (6) which provide similar solutions for working with annotations and LMD scopes. QuPath-to-LMD provides an easy-to-use application online and largely focuses on exporting QuPath annotations to LMD XML files. pyLMD provides exporting functionality, but with additional annotation management, and extensive documentation with examples. In the present work, we extend these efforts in our development of ShapeUpLMD, an open-source, Python-based solution that provides flexible control for LMD ROI optimization parameters, including shape, size, and spacing. ShapeUpLMD further introduces the ability to generate spatially resolved ROIs to support spatial multi-omics workflows. Together, these capabilities provide a comprehensive solution for transforming digital pathology annotations into regionally resolved, LMD-optimized coordinate files compatible with automated collection workflows.

We framed the performance of ShapeUpLMD in a cohort of USC tumors, leveraging expert pathologist annotations of tumor and non-tumor cell populations to train robust tumor cell-specific classifiers using the HALO digital pathology platform. Classifier performance was validated by direct comparison with manual annotations, demonstrating high concordance between manual and automated annotation strategies. Notably, these findings indicate that our workflow for training cellular subpopulation-specific ROI classifiers (Supplementary Figure 1) can be readily extended to other digital pathology platforms capable of performing automated classification and ROI export, such as QuPath (11). We further show that ROIs generated by third party software tools like HALO can be directly processed by ShapeUpLMD to generate optimized regions which yield high fidelity LMD collections, confirmed by our post-collection spatial concordance analysis. In summary, our results demonstrate that ShapeUpLMD-generated ROIs generated are highly concordant with manual annotation and, with our prior automation methods (2).

ShapeUpLMD also enables generation of spatially defined ROIs tailored to user defined tissue areas and parameters. In our proof-of-concept application, we subdivided a representative USC tumor into multiple spatially defined regions and performed global proteomic profiling. The data revealed distinct global protein profiles between whole tumor and enriched tumor regions. Proteins elevated in whole tumor regions included multiple collagen isoforms consistent with admixture of non-tumor cells (i.e. stroma cell populations) (1,2). The proteomic profiles from spatially resolved ROIs clustered according to local proximity, revealing region-specific proteomic consistency within tumors and underscoring heterogeneity across tumor cell populations. To further assess ShapeUpLMD for unbiased spatial proteomics, we generated evenly sized ∼1 mm² ROIs spanning an entire USC tumor tissue section. Tumor and non-tumor classifications derived from digital pathology strongly correlated with tumor and non-tumor cell admixture scores inferred directly from companion spatial proteomics data (7), confirming that spatially separated ROIs retain the unique protein abundance profiles common to tumor and non-tumor cell populations (1,2,9,10). These findings illustrate that ShapeUpLMD not only improves the precision of LMD-based enrichment but also facilitates spatially resolved proteomic analyses that uncover regional molecular variation otherwise masked in bulk tumor profiling.

Given the demonstrated recovery of high-quality DNA, RNA, and protein from LMD-harvested tissues (1), ROIs generated by ShapeUpLMD should prove useful for multi-omic workflows incorporating next-generation sequencing. Notably, ShapeUpLMD provides flexibility to users to define ROI area size and count, making it adaptable to diverse experimental designs and tissue types.

Limitations of this study include the modest cohort size and focus on a single malignancy (USC). ROI precision may vary across different LMD microscope platforms, laser parameters and imaging magnification. However, ShapeUpLMD is designed for broad adaptability through user-defined parameters, while also being readily refinable for diverse tissues and digital pathology ecosystems.

In summary, ShapeUpLMD provides an open-source solution for automated generation of LMD-optimized and regionally resolved ROIs. By bridging digital pathology, machine learning, and automated LMD, this pipeline enables reproducible, high-throughput, and spatially resolved tissue enrichment. As spatial multi-omics evolves toward comprehensive tissue atlases and clinically integrated diagnostics, ShapeUpLMD provides a scalable foundation for high-fidelity sampling across laboratories, diseases, and analytical platforms – advancing precision pathology and accelerating the next generation of spatial systems biology. ShapeUpLMD is available at: https://github.com/GYNCOE/ShapeUpLMD.

## Declarations

### Ethics approval and consent to participate

Fresh-frozen tumor tissues were selected from patients diagnosed with uterine serous carcinoma enrolled in a WCG IRB approved #20110222 Tissue and Data Acquisition Study of Gynecologic Disease who underwent surgery at Inova Fairfax Medical Campus (Falls Church, VA, USA), University of Oklahoma Health Sciences (Oklahoma City, OK, USA) or at University of Virginia (Charlottesville, VA, USA)

### Consent for publication

Not applicable

### Availability of data and materials

ShapeUpLMD is available at: https://github.com/GYNCOE/ShapeUpLMD; please contact for access. Representative input image for ShapeUpLMD testing is available here: https://lmdomics.org/ShapeUpLMD/. Proteomic data generated in this study is accessible within the Mass Spectrometry Interactive Virtual Environment (MassIVE) resource under accession MSV000100553 and the ProteomeXChange (PRIDE) resource under accession PXD073411.

## Authors’ contributions

J.S., D.M., T.P.C and N.W.B. lead the study. D.M. and S.T. performed identification of clinical specimens, D.M., K.N.W. and K.A.C., performed sample collections and molecular extraction. B.L.H, J.L., N.W.B., J.O. generated and analyzed proteomics data. P.M.F. performed pathology review. J.S., D.M., T.P.C and N.W.B wrote the manuscript, S.T., K.N.W., K.A.C., J.L., B.L.H, T.A., A.L.H., A. L. K.M.D., C.M.T., G.L.M reviewed the manuscript. All authors reviewed and approved the final manuscript.

## Acknowledgements

We would like to acknowledge Emily A. Pennington for her review and feedback on the ShapeUpLMD software code.

## Disclaimer

The views expressed herein are those of the authors and do not reflect the official policy of the Uniformed Services University of the Health Sciences, the Henry M. Jackson Foundation for the Advancement of Military Medicine, Inc., Inova Health System, the Department of Army/Navy/Air Force, Department of War, or U.S. Government. Mention of trade names, commercial products, or organizations does not imply endorsement by the U.S. Government.

## Figure Legends

**Supplementary Figure 1.**
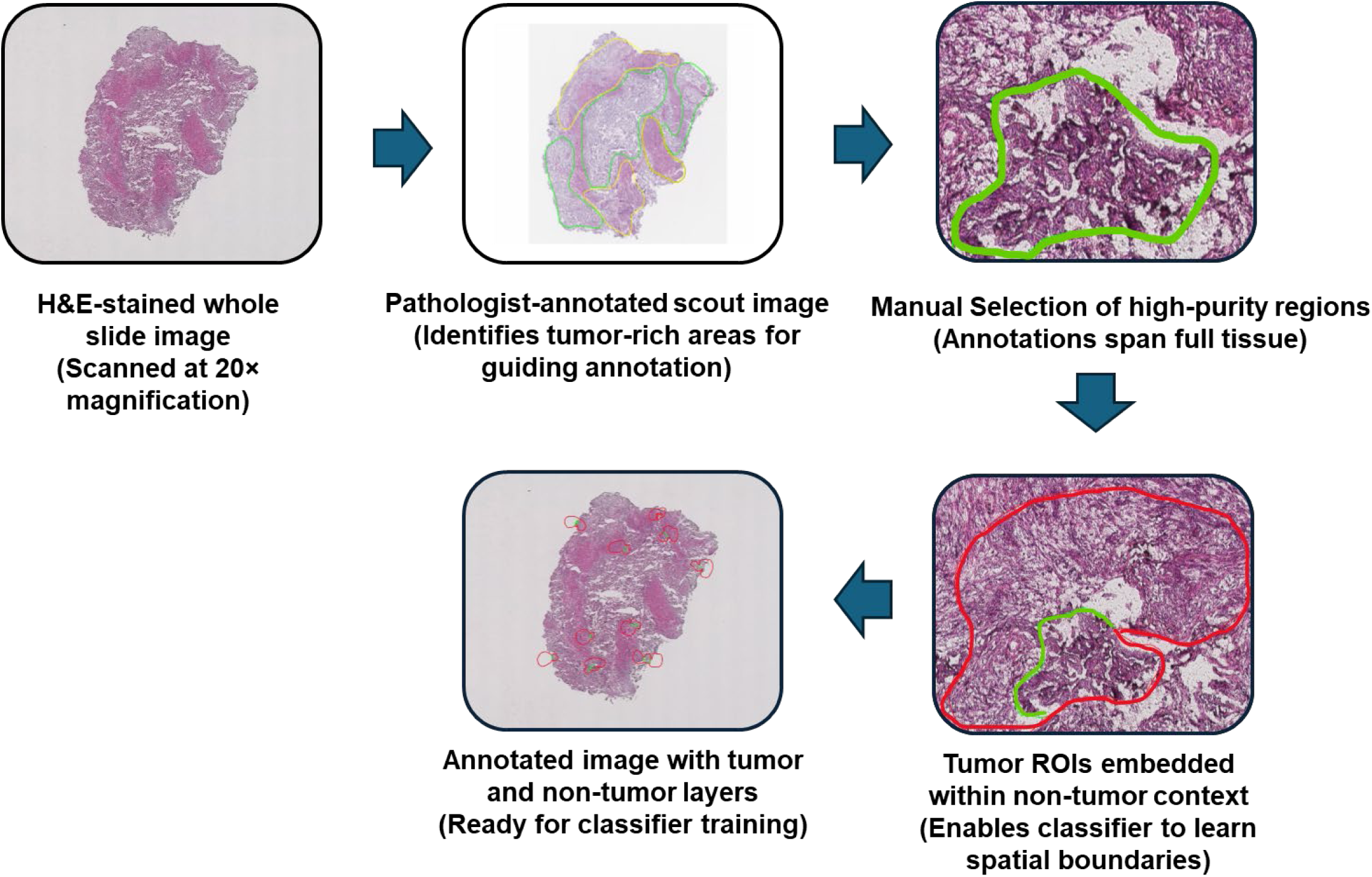
Workflow for generating automated classifiers of tumor cell populations in the HALO Image Analysis Platform.

**Supplementary Figure 2.**
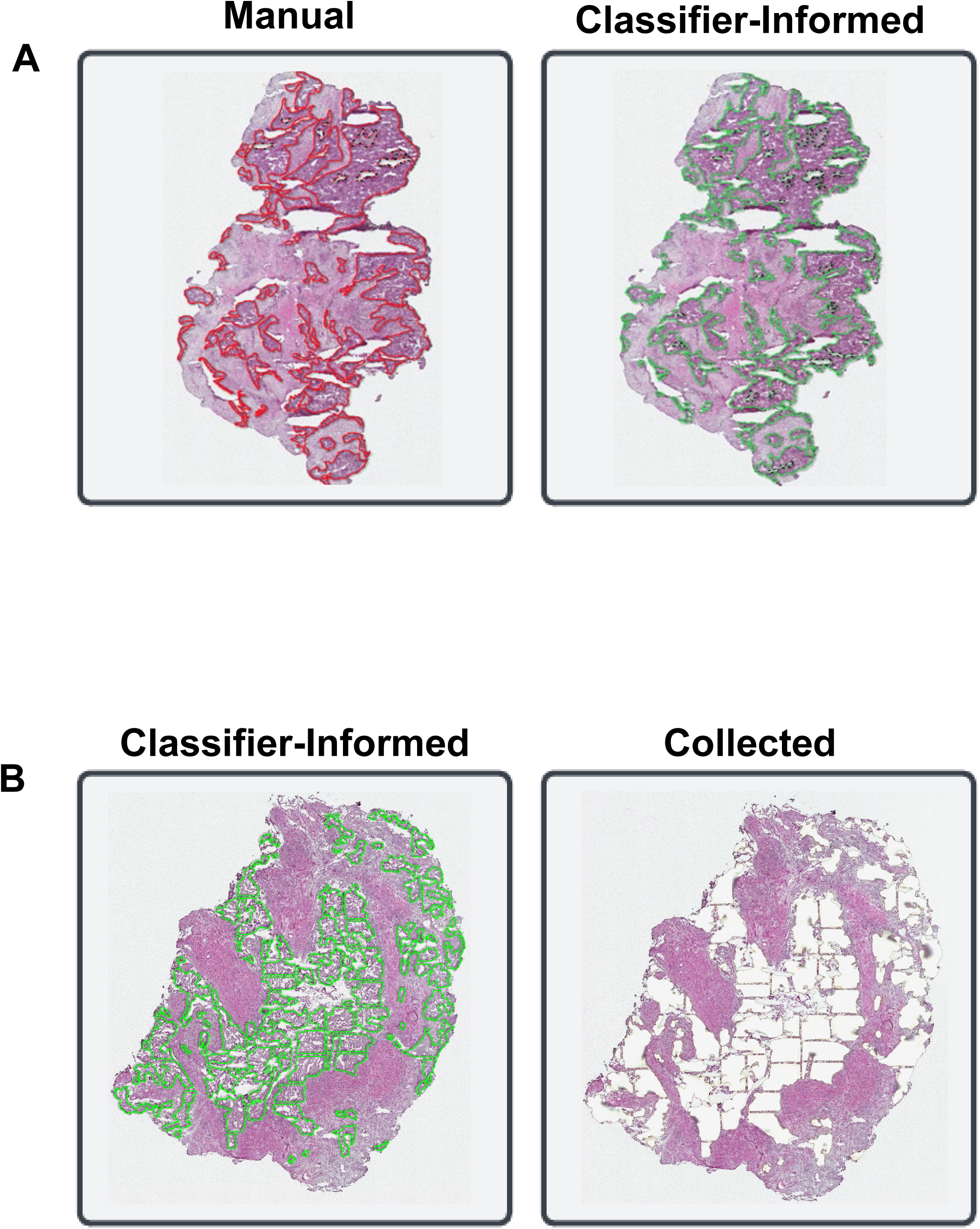
A: Representative USC tumor tissue section with matched tumor regions notated manually or using our automated classifier method. B: Representative USC tumor tissue section with matched tumor regions annotated using ShapeUpLMD-optimized tumor ROIs (classifier) or following collection of ROIs of interest by LMD.

**Supplementary Figure 3.**
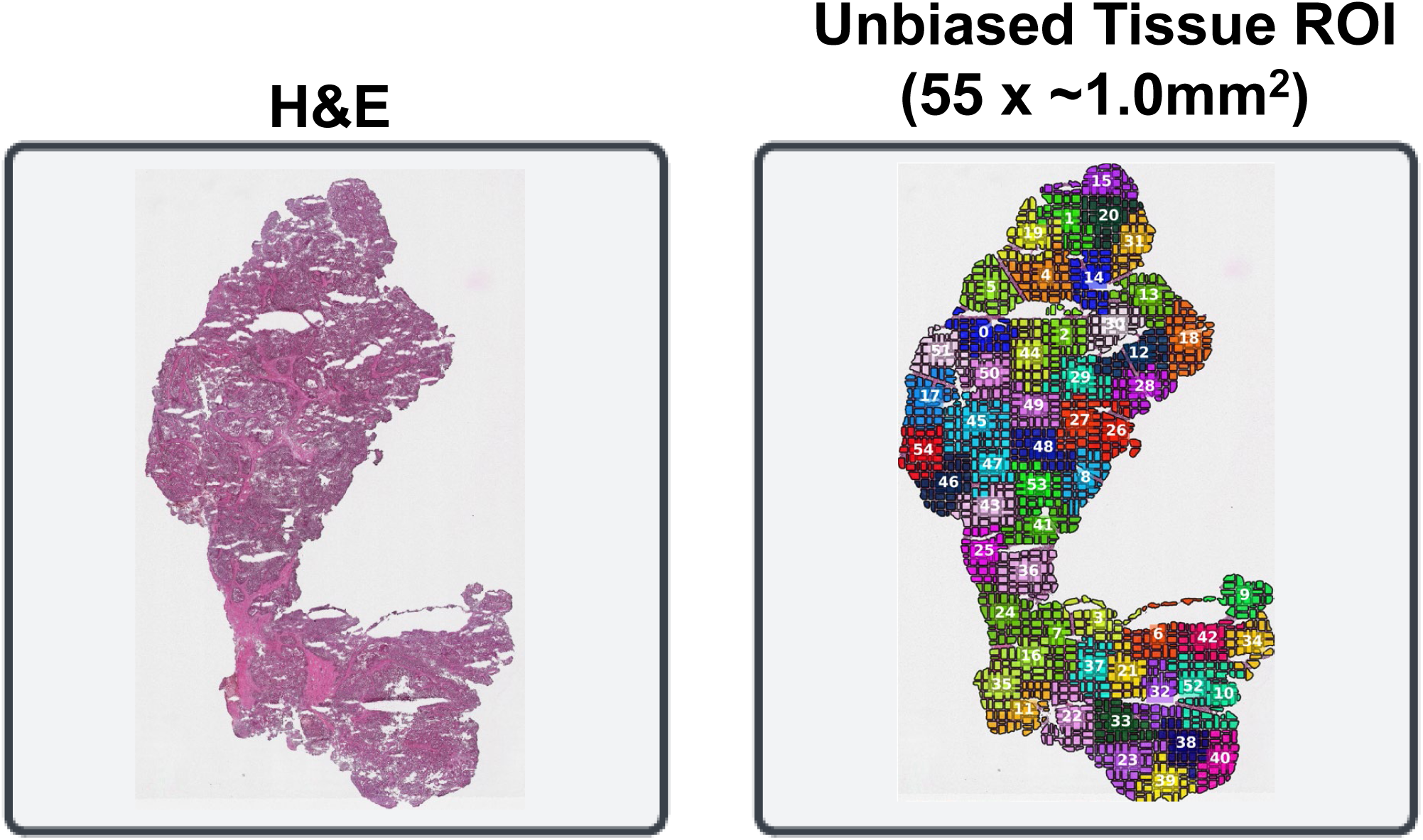
Representative USC tumor H&E stained tissue section and fifty-five ShapeUpLMD-optimized, unbiased tissue ROIs ± ∼1.0 mm^2^ spanning the entire tissue section.

## Supplementary Table Legends

**Supplementary Table 1:** Performance metrics comparing manual versus automated classifications of tumor ROIs for n11 USC tumor tissue samples.

**Supplementary Table 2:** Performance metrics comparing estimated collected areas versus automated classifications of tumor ROIs for n11 USC tumor tissue samples.

**Supplementary Table 3:** Z-test results comparing proteome alterations in average enriched tumor versus whole regions.

**Supplementary Table 4:** Pathway analysis of proteins elevated in enriched tumor versus whole tissue collections.

**Supplementary Table 5:** Tissue section ROIs for unbiased spatial proteomics analyses; details tissue regions, ROI area (mm^2^) as well as percent tumor, stroma or other admixture predicted from automated classifiers generated using the workflow in Supplementary Figure 1 informed from review of companion H&E tissue sections.

**Supplementary Table 6:** ProteoMixture scores calculated from spatial proteomics data and CDH1, EPCAM, FAP and COL1A1 protein abundance from unbiased spatial proteomics analyses.

